# *SIMPROV* : Provenance capturing for simulation studies

**DOI:** 10.1101/2025.02.26.640288

**Authors:** Andreas Ruscheinski, Anja Wolpers, Philipp Henning, Pia Wilsdorf, Adelinde M. Uhrmacher

## Abstract

Improving interpretability and reusability has become paramount for modeling and simulation studies. Provenance, which encompasses information about the entities, activities, and agents involved in producing a model, experiment, or data, is pivotal in achieving this goal. However, capturing provenance in simulation studies presents a tremendous challenge due to the diverse software systems employed by modelers and the various entities and activities to be considered. Existing methods only automatically capture partial provenance from individual software systems, leaving gaps in the overall story of a simulation study. To address this limitation, we introduce a lightweight method that can record the provenance of complete simulation studies by monitoring the modeler in their familiar yet heterogeneous work environment, posing as few restrictions as possible. The approach emphasizes a clear separation of concerns between provenance capturers, which collect data from the diverse software systems used, and a provenance builder, which assembles this information into a coherent provenance graph. Furthermore, we provide a web interface that enables modelers to enhance and explore their provenance graphs. We showcase the practicality of *SIMPROV* through two cell biological case studies.

**Author summary:** With the importance of simulation studies in understanding and managing complex dynamic systems, the need to support the interpretation and (re-)use of their results increases. Provenance documents how the products of a simulation study were created and what other products, agents, and activities have been involved in this process. For example, the information based on which data from which cell line a simulation model has been calibrated and validated is central to interpreting the results and assessing how the results can be reused. Therefore, some software tools offer to record provenance information. However, for complete provenance information, the tool must offer all functionalities required for a simulation study. In practice, various tools are typically used. To accommodate this situation, we propose a flexible, decentralized approach: *SIMPROV*. A provenance capturer – a small piece of software designed to record the modeler’s actions within a software tool – observes each tool used by the modeler. A central provenance builder then combines the recorded information from all captures. A capturer has to be programmed only once for each software tool used in systems biology, and modelers can work as before with minimal effort needed to record the provenance of their simulation studies automatically.

## Motivation

Established reporting and documentation guidelines in systems biology enlist information required for interpreting and reusing simulation models [1] and simulation experiments [2]. The former includes assumptions and decisions made during model construction [3] and semantic annotations for relating model parts to real-world entities [4]. Reporting guidelines such as TRACE [5] and STRESS [6] direct the attention towards the entire simulation study and document, e.g., calibration and validation steps that have been conducted. Similarly to the COMBINE archive [7], they bundle heterogeneous information sources that are relevant in reproducing and interpreting the results of a simulation study. However, neither captures the processes by which activities, information sources, and the generated artifacts are interrelated. Simulation studies comprise iterative processes of collecting information about the system of interest, specifying and refining the simulation model, checking the expected behavior of the simulation model, conducting simulation experiments, and analyzing the results. Thus, these documentation guidelines leave significant gaps in the overall story of a simulation study even though the documentation of all steps has been identified as crucial for credible simulation models [8] and for justifying the decisions made during the simulation study [3].

Provenance embraces any information describing the production process of a product of interest [9]. In particular, it refers to “information about entities, activities, and people involved in producing a piece of data or thing” [10]. Thus, provenance moves the simulation study processes into the focus of interest. It puts the different entities of a simulation study, e.g., assumptions, requirements, data, simulation experiment specifications [7], and different versions [11] and components [12] of simulation models into a specific context [13], i.e., how they have been used and generated during the simulation study. Thus, provenance can enrich existing collections of artifacts such as COMBINE archives [7]. With provenance, not only do the final products become accessible, but intermediate products and how they relate to each other, complementing existing simulation model documentation formats [14–16]. Moreover, provenance has been found useful for relating different simulation studies to become a family of models [17], and for documenting the process of pursuing different hypotheses to finally arrive at a simulation model that fits the findings of different wet-lab studies [18] (see Fig. 1). Thus, provenance appears crucial for improving transparency, reproducibility, and reuse of simulation models and data from simulation studies [4, 5, 19–21], and for automatically executing simulation studies and the simulation experiments therein [22, 23].

**Fig 1.**
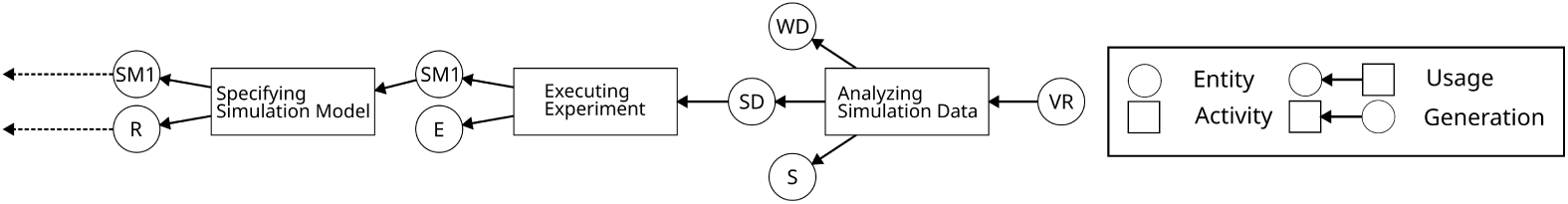
Excerpt from a provenance graph depicting the refinement process of a simulation model (SM1) during a simulation study. This includes the incorporation of references (R) in the refinement, followed by testing the model’s ability to reproduce wet-lab data (WD) through a simulation experiment (E) in which the resulting simulation data (SD) is then analyzed using a Python script (S) to obtain the validation result (VR).

However, manually recording the provenance of a simulation study will, in most cases, be unrealistic due to the implied effort, also it bears the danger that incomplete or incorrect information might enter the documentation [24]. The obvious solution is to gather as much provenance information as possible while conducting the simulation study automatically and in a structured manner.

Workflows are typically equipped with means of automatically recording provenance [25]. However, simulation studies belong to a class of processes that are not well supported by established workflow management systems, as these knowledge-intensive processes require substantial flexibility at both design- and run-time [26]. Declarative workflows offer the required flexibility and support [27] and have been used for conducting simulation studies [28] and for the automatic recording of provenance information [29]. However, to use the automatic documentation of workflow systems, the modeling and simulation study has to be conducted within these workflow systems, which need to support the specific combination of tools of relevance for the simulation study being conducted [8].

Here, we will opt for provenance documentation that avoids the additional workflow layer. Instead, our approach provides a modular architecture that records the modeler’s activities in each tool used for the simulation study separately via *provenance capturers* and then centrally collects, interprets, and aggregates the provenance information in a *provenance builder*.

In the following, we will first overview the W3C provenance standard PROV-DM and describe how it can represent and store provenance information about a simulation study. Afterward, we will introduce the architecture of our lightweight and modular provenance-capturing approach and the methods involved and describe its implementation in the web-based tool *SIMPROV*. Next, we will demonstrate *SIMPROV* in two cell biological case studies. In the first case study, we capture developing and extending an mRNA delivery model [30] within a new modeling and simulation tool for rule-based modeling with dynamic compartments [31]. In the second case study, we used Tellurium [32] to replay crucial steps of a simulation study that analyzed endocytosis processes of the Wnt signaling pathway [18] and compared the generated provenance graph to the one that was manually generated and included in the original publication. We close the paper with a discussion, conclusions, and future work.

## Provenance of Simulation Studies

Recording the provenance of any process requires a *provenance model* prescribing how the recorded provenance information is formatted. In addition to *how* the provenance information is formatted, the choice of the model also defines *what* information is needed to form said structure: It dictates the minimum of information needed and what optional information can be included [33, 34].

### PROV-DM Extension for Simulation Studies

Our provenance model is based on the W3C provenance standard, PROV-DM [35], as PROV-DM has previously been extended to develop a provenance model for entire simulation studies [15, 17, 36, 37]. In PROV-DM, the provenance information forms a directed acyclic graph in which the nodes represent *entities*, *activities*, and *agents*. The edges are used to describe their *dependencies*, i.e., the following relations:

1. *usage-relations* (entity *←* activity) that describe which entities have been used in the activity,
2. *generation-relations* (activity *←* entity) that describe which entities have been generated by an activity and
3. *association-relations* (agent *←* activity) that indicate an agent’s responsibility for an activity that took place.

We extended PROV-DM with *entity types* representing a study’s intermediate and final products and *activity* types describing what activities are executed by a modeler [15, 17, 36], which will represent the provenance model applied by *SIMPROV*. What *activities* are executed by a modeler and what *entities* are part of these activities throughout a simulation study has been identified in various life cycles [29, 38, 39] and reporting guidelines [5, 6] of simulation studies. According to those, a simulation study starts with a *research question* about a specific aspect of a real-world system that shall be answered by modeling and simulation. Next, a *conceptual model* (which is often said to include the research question [40]) is built that contains all context that informs and guides the model building process including *requirements*, *assumptions*, a *qualitative model*, and further *data* and *information* [38, 41]. Based on the conceptual model, the *simulation model* can be implemented, e.g., via a domain-specific modeling language, which can then be executed as part of a *simulation experiment* to produce *simulation data*. Until the research question can be answered based on this data, the simulation model is successively refined, and the new model versions are calibrated, validated, and analyzed by simulation experiments, as represented by *activities*. Additionally, various *postprocessing* steps of the experiment outputs may be necessary, producing entities such as *scripts* or *visualizations*. The software systems used by the modeler during these activities are represented as *agents*. According to PROV-DM, agents refer to anything that bears responsibility for generated entities and activities that happened. Therefore, we propose that the software systems used by the modeler during these activities are represented as *agents*. These may be the simulators used to run the model or the software tools used by the modeler to analyze and visualize the experiment results. Treating software as agents (rather than meta-data of other entities, such as simulation experiment specifications) underlines the importance of software in simulation studies for reproducing simulation results.

Table 1 provides an overview of the central entities in a simulation study. In the following, we discuss in which formats they typically are provided and their role in simulation studies.

**Table 1.** Provenance entities of a simulation study.

| Entity | Form |
| --- | --- |
| **Conceptual Model** |  |
| Research Objective | text, any media type |
| Requirement | reference to data, logic formula, domain-specific language |
| Assumption | text, formula |
| Qualitative Model | text, sketches, diagrammatic formalism |
| Data | CVS, JSON, XML, spreadsheets, etc. |
| Reference | publication, any media type |
| **Simulation Model** | general-purpose language,<br>modeling formalism,<br>domain-specific language |
| **Simulation Experiment** | general-purpose language,<br>domain-specific language |
| **Simulation Data** | CVS, JSON, XML, spreadsheets, etc. |
| **Postprocessing** |  |
| Script | general-purpose language,<br>domain-specific language,<br>scientific workflow |
| Visualization | any visual media type |

#### The Conceptual Model

The importance of the conceptual model in designing a simulation model and conducting a simulation study is widely acknowledged [38, 39, 41]. Although the interpretation of the conceptual model varies [40], one widely accepted definition is that the conceptual model is a collection of all types of resources that inform the construction of the simulation model [38, 39]. Thus, the conceptual model refers to early-stage products of a simulation study [42]. In our data model, we include the following entities as, e.g., listed by [17, 40].

*Research Questions* define what questions or objectives are sought to be addressed by the study. They determine the focus in constructing the model and which experiments have to be executed to reach answers. Thus, it is essential to include the research questions in recording provenance, as also highlighted by reporting guidelines [5].

*Requirements* define the simulation model’s output behavior and further substantiate the research question. Throughout the simulation study, the model is checked and refined to ensure that the requirements are satisfied via a variety of calibration and validation activities [40]. Therefore, requirements play a crucial role in motivating the specification and execution of simulation experiments as part of provenance. They may be encoded by a reference to data that shall be reproduced [40], by a logic formula (such as Metric-Interval Temporal Logic [43]), or in a domain-specific language for requirement specification (e.g., the Biological Property Specification Language [44]).

*Assumptions* describe the scope of a model and what simplifications were made throughout the simulation study to reduce the level of complexity. Therefore, documenting assumptions is needed to correctly build the model and to interpret the simulation study’s results. Assumptions can be formulated using arbitrary formulas with a defined mathematical syntax, or free text. Different types of assumptions may be annotated with an ontology, such as the Systems Biology Ontology (SBO) [45], to indicate which part of the model contains which kinds of assumptions [17]. For example, the assumption that the concentration of a specific model entity remains constant during the simulation refers to SBO term 362 (concentration conservation law) [17].

The *Qualitative Model* is an early, often more abstract version of the simulation model focusing on causal relationships. Qualitative models can be specified using various formats, such as informal sketches and textual descriptions of varying complexity, that can help design the model. But also qualitative formalism with a visual (diagrammatic) representation may be used, such as causal loop diagrams [46] or the Systems Biology Graphical Notation [47].

The final element of the conceptual model is *data* used as input for variables and parameters or used for calibration or validation.

All these elements might be subject to further substantiation by *references* to scientific papers, websites, or other source types.

#### Simulation Models

Based on the entities in the conceptual model, a simulation model is specified [38, 39]. A (simulation) model “(M) for a system (S) and an experiment (E) is anything to which E can be applied in order to answer questions about (S)” [48]. So, in contrast to the conceptual model, the simulation model is executable by a simulation experiment via a simulator. A simulation model may be implemented using a modeling formalism like (stochastic) Petri nets [49], or using domain-specific languages [50] like rule-based languages, e.g., the BionetGen language [51], the Systems Biology Markup Language (SBML) [52], or CellML [53]. In addition, general-purpose programming languages (e.g., Python or Julia) may be used to implement the model.

#### Simulation Experiments

A simulation experiment is “the process of extracting data from a system by exerting it through its inputs” [48]. They may be specified using domain-specific languages for experiment design, such as SESSL [54] or SED-ML [55], based on meta-models [23, 96] or in a general-purpose language, such as Python or R, with the option to use dedicated libraries for experiment design [56]. Various types of simulation experiments are specified and executed with different goals throughout a simulation study, e.g., to calibrate, validate, and analyze the simulation model [39, 57]. For instance, a modeler may aim to use sensitivity analysis to probe a model’s robustness [58], compare simulation data of related models for cross-validation [59], or check existing hypotheses regarding a biochemical mechanism using parameter scans [18].

#### Simulation Data

Executing simulation experiments produces *simulation data* that can then, e.g., be compared to reference data from the conceptual model for calibration or validation. Furthermore, simulation data may be reused in other simulation studies and thus become part of the conceptual model (in those subsequent studies). The format of simulation data depends on the tool running the simulation experiment and the simulator used.

#### Postprocessing

Several postprocessing steps may be required depending on the answers the modeler wishes to extract from simulation data. This includes specifying and executing scripts to transform data and produce visualizations.

*Scripts* are additional executable files, that are crucial for extracting relevant insights from given simulation data. Scripts may be specified in any general-purpose programming language, in a domain-specific language for creating visualizations [60], or in a scientific workflow format [61].

*Visualizations:* The appropriate type of visualization depends on the experiment executed, the data produced, and the information the modeler wants to convey. Thus, in general, any visual media format may be used.

Figure 2 shows a provenance graph in the graphical notation of PROV-DM, illustrating various types of entities, activities, agents, and dependencies discussed above.

**Fig 2.**
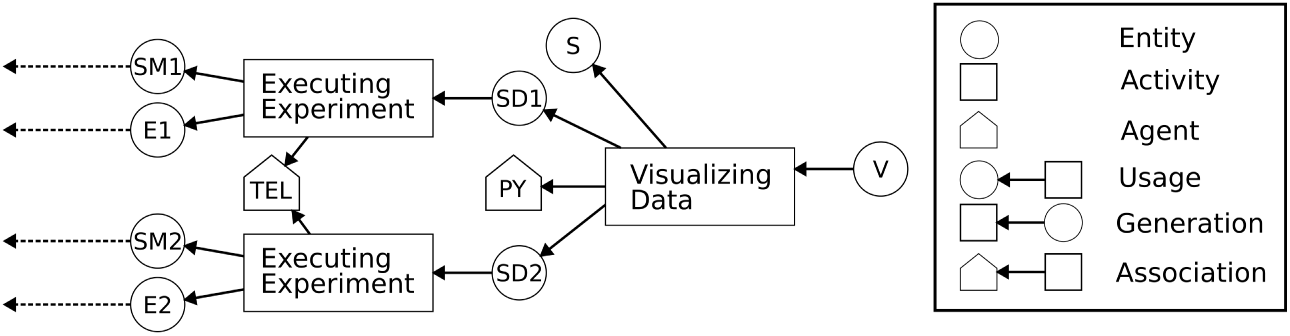
Example provenance graph of exploring the difference between two simulation models by visualizing simulation data generated from two independent simulation experiments. The *activities* comprise the executions of the experiments (Executing Experiment) and the visualization of results (Visualizing Data). The *entities* comprise the two simulation models (SM1, SM2), two simulation experiments (E1, E2), the simulation data generated by the experiments (SD1, SD2), and the Python script (S) for generating visualization of the simulation data (V). The *agents* are the modeling and simulation tool (TEL) and the programming environment (PY).

### Provenance Patterns

Based on the extension of PROV-DM for simulation studies we formulated *provenance patterns* [22] that define which combinations of provenance entities, dependencies, and agents are valid for an activity type.

For example, the activity “Specifying Experiment” always uses a simulation model entity but can sometimes use an assumption entity and a requirement entity (see Fig. 3). In addition, a “Specifying Experiment” activity always produces a simulation experiment entity. In addition, we defined required and optional attributes for the entities and agents as part of the *provenance patterns*.

**Fig 3.**
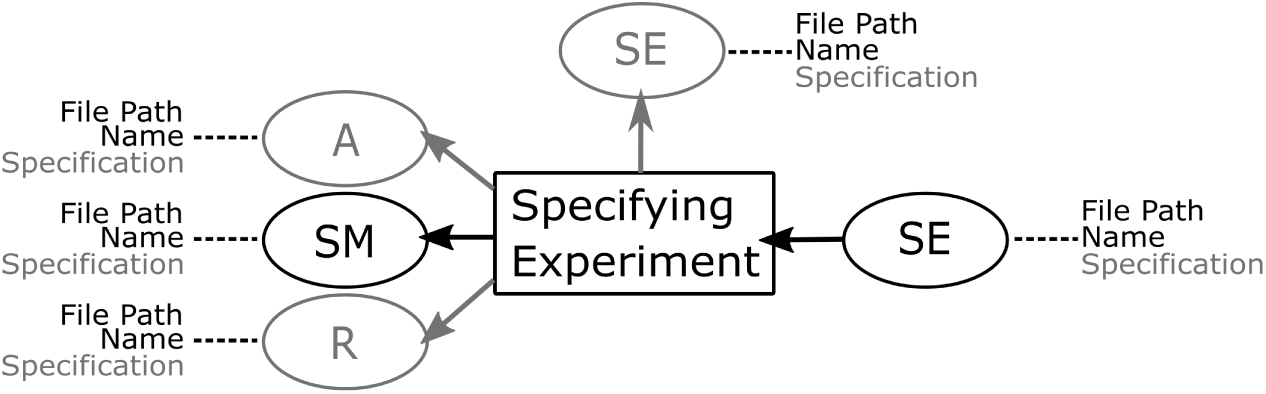
An example for a provenance pattern for the activity *Specifying Experiment*. The activity always uses a simulation model (SM). Optionally (denoted in gray), it can also use an already existing simulation experiment, an assumption (A), and/or a requirement (R). The attributes required or optional (gray) for the entities and agents are annotated next to the dashed lines. So, if an activity uses a requirement, then the requirement must have attribute values for its name and file path – its specification can be blank.

Based on the life cycles of simulation studies and guidelines discussed above, we specified patterns for the activities *Specifying Reference*, *Specifying Research Question*, *Specifying Requirement*, *Specifying Assumption*, *Specifying Simulation Model*, *Specifying Simulation Experiment*, *Executing Simulation Experiment*, and *Analyzing Simulation Data*.

## Concept

To collect provenance information from a varying set of individual tools, we propose a modular architecture that can be adapted to the modeler’s needs for different tool setups. We adopt Eek and Miller’s idea [62] to capture provenance from individual tools locally before combining all captured information centrally. They proposed an “Agent-Based Provenance Architecture” [62] introducing various types of agents to capture provenance from distributed, heterogeneous software applications. Here, we propose an architecture tailored to record the provenance of simulation studies.

Our approach is highly flexible and can be easily adapted to the individual modeler’s needs. It consists of two module types: provenance capturers and a (central) provenance builder (see Fig. 4). The *provenance capturers* are responsible for detecting and collecting information about the modeler’s activities from the different software systems or tools. As modelers use a wide range of systems, tools and their combinations they can combine any number of capturers – one captuer for each system or tool. Whenever a capturer detects that the modeler performed an activity using the capturer’s software system or tool, it collects all relevant information including which entities and agents are involved in the activity and communicates it to the *provenance builder*.

**Fig 4.**
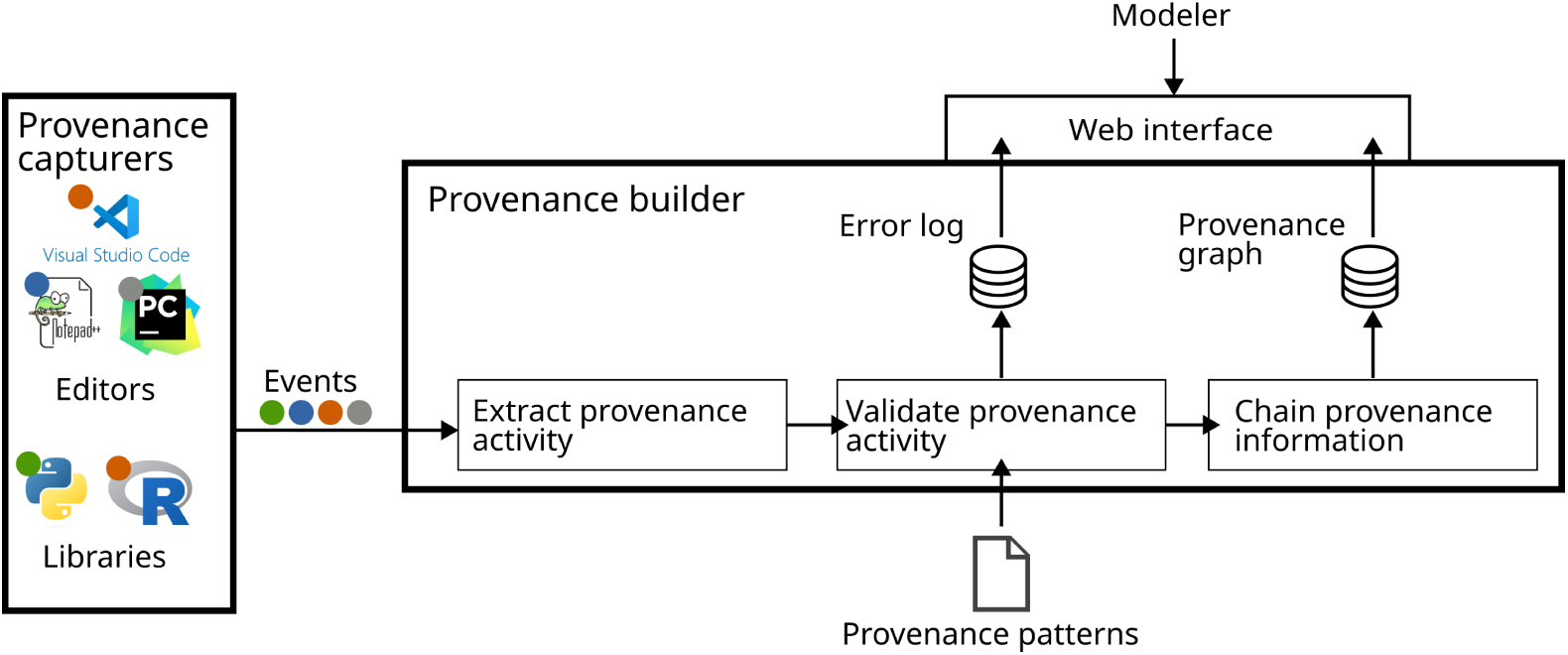
Overview of the *SIMPROV* software architecture. Provenance capturers record information on the tools used by the modeler. The provenance builder collects and combines the captured information into a provenance graph.

The *provenance builder* processes the incoming events and checks if the new provenance activity with its entities and the associated agents matches the provenance model (incl. the provenance patterns) used by the simulation study. Here, the provenance model (incl. the provenance patterns) will be as discussed above. Still, other provenance models can also be used with our approach (you only have to adapt the provenance capturers accordingly). If the new activity, including its entities and agents, matches the activity’s provenance pattern, it is chained with the already captured provenance graph.

### Provenance Capturers

A *provenance capturer* has to accomplish four tasks:

1. Detecting the modeler’s activity in a software system,
2. Collecting information about the activity,
3. Identifying the activity and related entity types, and
4. Sending the information to the *provenance builder*.

#### Detecting the modeler’s activity & Collecting information

Given the diverse software architectures used in various systems, the methods by which a *provenance capturer* detects and collects information about the modeler’s activity must be determined case by case, depending on the type of software observed by the respective capturer. Previous studies employ numerous approaches to capturing provenance from computational tasks ranging from observing the modeler’s operating system on the level of system calls [63] to providing an annotation syntax that modelers can include in their code [64]. The information gained from recording system calls is recorded completely automatically, but it also includes information irrelevant to the simulation study that is complicated to eliminate. On the other hand, asking the modeler to manually annotate their code requires continuous additional effort on their part, which we aim to minimize. So, we recommend using methods that compromise between recording targeted information and user involvement. We argue that two methods, using plugin interfaces and wrapping library functions, suffice to capture provenance from most types of software.

*Using plugin interfaces* We suggest using the readily available plugin interfaces for editors and IDEs, such as PyCharm [65], Visual Studio Code [66], or Notepad++ [67]. These plugin interfaces typically define functions executed whenever the underlying software system *detects* a user interaction, such as clicking a button or saving a file. In this case, the provenance capturer consists of a custom function that introspects the information about the user interaction to *collect* information about the modeler’s activity, which will then be used to *identify* the activity and the associated entities and agents.

*Wrapping library functions* When using software libraries, such as Hybrid Automata [68], Matplotlib [69], Tellurium [70] or SESSL [54], the modeler calls the library functions to execute tasks like running a simulation or analyzing the simulation data. Monitoring the called functions can *detect* the modeler’s activity in this context. Thus, we can implement a provenance capturer via wrapper functions that call the original function and introspect the function arguments and results to *collect* information about the activity.

Further, one-stop simulation systems, such as COPASI [71] or NetLogo [72], that provide an IDE for specific simulation models and use a simulation library for running simulation experiments with them might require combining both capturing approaches.

#### Identifying the activity and entities

Independently of whether provenance has been captured based on an IDE or libraries, we need to identify the activity and entity types.

*Entities:* First, the provenance capturer identifies the type of all related entities. In general, the entities’ types can be deduced from the file’s content, its name (extension), or other conventions that make the file’s entity type explicit. Ideally, the provenance capturer can deduce the file’s entity type from its content or name. However, this is not always possible and the modeler has to make the entity type explicit – for example, by adhering to naming conventions. Which of (combinations of) these approaches can be used differs for the different entity types and depends on the modeler’s work practices.

Naturally, the interpretation of a simulation experiment or simulation model specification is facilitated by accessible syntax and semantics, e.g., in terms of meta-models about simulation experiments [23] or a formalization of a model language [73]. This also applies to formally specified behavioral requirements to be checked automatically [44, 74]. In addition, such structures enable the *provenance capturer* to recognize other entities (from the conceptual model) used by the simulation model or experiment. For example, an experiment might import a behavioral requirement as a logic formula to check if the formula holds for a simulation model (the produced simulation data) in a certain configuration. As the behavioral requirement is directly imported, its involvement in the experiment execution can be inferred by inspecting the experiment file for imports.

Now, suppose the behavioral requirement was only available in text form. In that case, the experiment cannot import it directly. Hence, the *provenance capturer* will have no way of knowing that a behavioral requirement was involved with the execution of the experiment. Moreover, the capturer will not be able to distinguish the behavioral requirement from an assumption or qualitative model, for example, as all these entity types can come in the form of text which is hard to interpret automatically.

These problems exist not only for files of the entity type requirement but for all elements of the conceptual model. Table 1 shows that no entity type from the conceptual model can be identified universally and distinctively just from their file format (e.g., both research questions and assumptions are commonly specified in plain text files). In addition, most entities in the conceptual model lack not only a formalized but also a standardized syntax for seamless integration into files used by other entities. For instance, even if a requirement is specified using a formal approach, such as a logical formula, it may not be directly usable by the simulation experiment, depending on the compatibility between the used requirement checking approach and the specification language. Thus, the provenance capturer cannot automatically detect that these entities were used in the creation or execution of said files.

Therefore, we still need the modeler to make this information explicit – in the form of naming conventions in file names or annotations in files, for example.

*Activity:* Based on the identified entities and the action performed by the modeler, the *provenance capturer* infers the activity’s type. The entity’s type has been inferred from the entity’s file name or file introspection as described above. The activity performed by the modeler can be identified from the function triggering the provenance capturer: when a file is saved, the activity is *specifying*, and when a file is executed, the activity is *executing*.

#### Sending the information

Finally, the *provenance capturer* encodes the provenance activity and entities in an *event* and sends it to the *provenance builder*. The *event’s* type corresponds to the activity type inferred in the preceding step. It includes information about all related agents and entities, including their attributes, such as file path, name, version, and content.

### Provenance Builder

As soon as the *provenance builder* receives a new event, it extracts the provenance activity with its entities and agents from the event. The *provenance builder* then has to accomplish two tasks:

1. Ensuring the consistency of the provenance information as defined by the provenance model and
2. Chaining the extracted provenance activities into a single provenance graph.

#### Ensuring Consistency

After unpacking the provenance activity with its entities and agents from the event, the *provenance builder* checks if the newly created provenance is complete and valid according to our provenance model. Using the defined *provenance patterns*, the model builder can not only check if all required entities are attached to the activity but can also make sure that all attributes such as file paths and names are available for each entity and agent.

If an attribute value is missing or invalid entities/agents are assigned to an activity, e.g., due to a lack of information in the original event, the activity is discarded and added to an error log that the user can inspect. Otherwise, the provenance activity (with its entities, agents, and dependencies) is chained with the already captured provenance graph.

#### Chaining into a Single Provenance Graph

Finally, the identified activity with its dependencies and entities is inserted into the provenance graph. The activity can be inserted directly because the activity has just been created and will not already exist within the graph. However, an entity can already be part of the provenance graph because the product the entity represents has been used or produced by a previous activity. Consequently, before inserting an entity, the application must check whether the entity to be inserted is already part of the graph. If this is the case, the entity is not inserted again, but the dependency is added between the latest version of the existing entity and the new activity. This is possible because whenever an entity is edited a new version of it is created as part of the recording.

## Implementation

We implemented *SIMPROV* adopting a client-server architecture in which *provenance capturers* function as clients and a *provenance builder* operates as the server (see Figure 4). As the provenance capturers have to be developed individually for the specific software system or tool used by a modeler, we will explain our provenance capturer’s implementation in our case study, below. Independently of the individual provenance capturer’s implementation, though, all provenance capturers send their captured information to the provenance builder as an *event* using arbitrary key-value pairs in JSON [75]. Afterward, the *event* has to be sent via a POST request to the REST API of the *provenance builder*.

The *provenance builder*, in contrast, stays the same independently of the individual modeler’s choices. It processes incoming events by extracting the captured information and converting it into objects representing provenance activities, entities, and agents connected with dependencies as defined in PROV-DM. This new excerpt of a provenance graph is then matched to the activity type’s *provenance pattern* that is declaratively specified using YAML [76]. For all recorded activities that could not be successfully matched with the respective provenance pattern, we keep an error log collecting the respective events for manual inspection.

Also, we equipped the *provenance builder* with a web interface allowing the modeler to explore the captured provenance graph visually, introspect the error log, and download the stored provenance information for further processing or publishing [77] (see Figure 5). For this, a Node.js app was implemented that communicates with the provenance builder via its REST API. In summary, the app allows the modeler to

1. visually explore the provenance graph rendered using Cytoscape [78],
2. download the provenance information in the PROV-JSON format [77] for reuse in other applications,
3. fill missing entity/agent attributes or add new dependencies between activities and entities/agents in case some information was not extracted by the rules, and
4. access the error log and events for debugging purposes.

**Fig 5.**
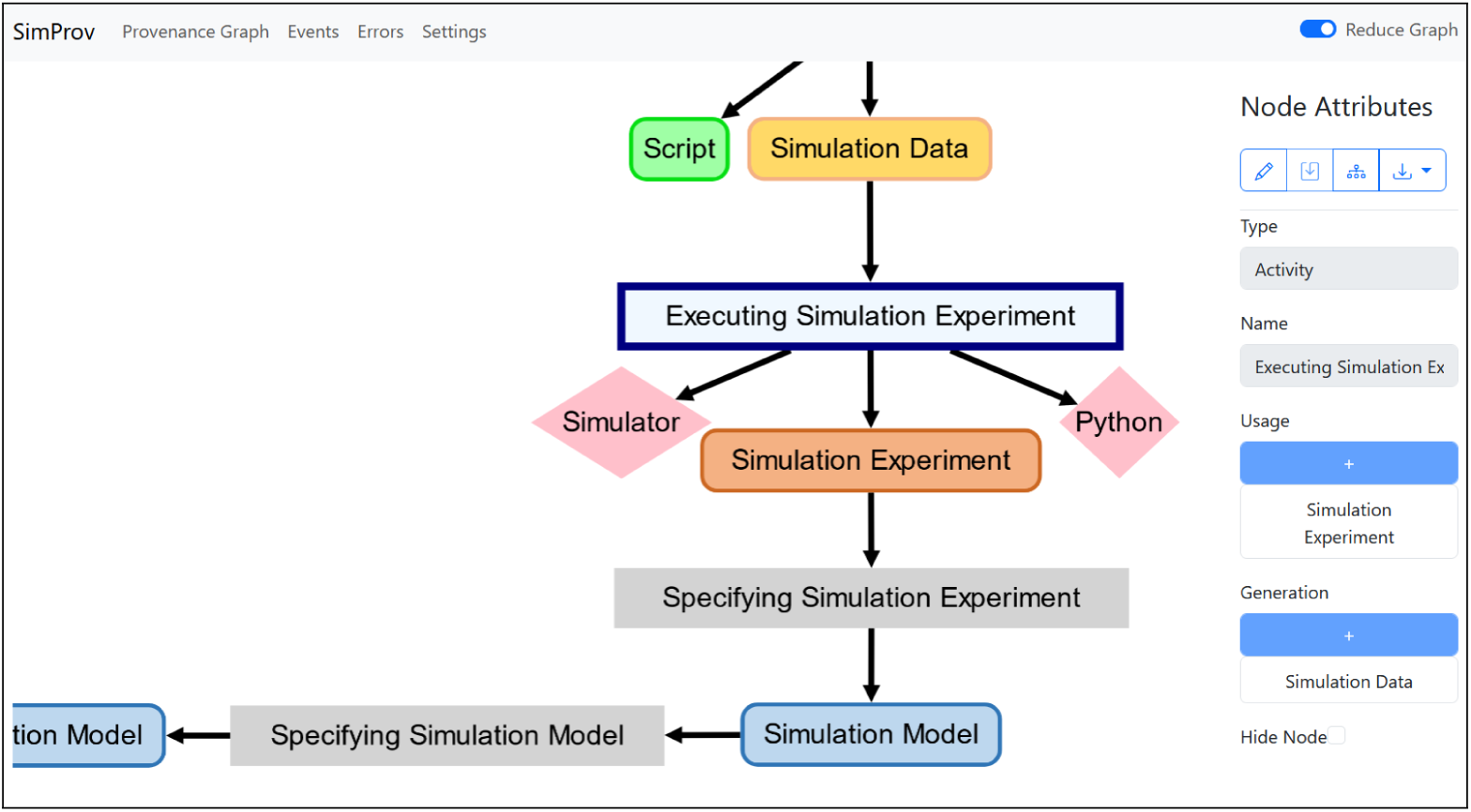
Screenshot of the *SIMPROV* web interface

Moreover, we also integrated various filtering approaches to reduce the number of nodes and edges shown in the graph, e.g., a custom abstraction-based filtering approach [29] that groups sequences of the same activity into a single activity, or a mechanism using the transitive reduction [79] of the provenance graph to remove edges.

## Case Studies

For the case studies, we reimplemented two models. First, we transferred and extended an mRNA delivery model from COPASI [71] to ML-Rules 3 [31], a simulation tool to efficiently simulate multi-level models including dynamic compartments. Based on this case study, we will explain the process of capturing the provenance graph in detail. Second, we took a Wnt/*β*-catenin signaling model originally implemented in ML-Rules 2 [73] and reimplemented it in Tellurium [32]. The original paper [18] includes a manually drawn provenance graph, which we will compare with the automatically captured *SIMPROV* graph.

### Capturing Provenance from Visual Studio Code and Python

For both of the following case studies, the modeler uses Visual Studio Code as an editor for specifying

1. the conceptual model, consisting of the research question, literature references, requirements, and assumptions, using Markdown [81],
2. the simulation models using the rule-based multilevel modeling language ML-Rules [31] or Tellurium, and
3. the simulation experiments and scripts for visualizing and analyzing simulation data using Python 3.10 (https://www.python.org/).

Note that *SIMPROV* is designed to work with any software or tool. So, the following only reflects how *SIMPROV* can work for the combination of software used in our case studies. One might even choose to implement the provenance capturer for the same software differently if it better matches the software’s internal workings.

As the modeler conducts the whole simulation study with Visual Studio Code, we implemented only one *provenance capturer*. The capturer employs two capturing methods in combination: using a plugin interface and providing the Python utility library “simprovhelper”.

The plugin makes use of Visual Studio’s command palette. The command palette offers a list of commands that work similarly to macros. A command can, for example, create a new file and ask the user to enter a name for it. Our capturer plugin provides commands to create files for each kind of provenance entity explained above (i.e., requirements, simulation models, simulation experiments, etc.). Depending on the entity type, the modeler is only asked to enter a name for the new file. For some entity types, the modeler might be asked, in addition, to select already existing entities from the conceptual model that are used to specify the new provenance product. For example, for the specification of a simulation model, assumptions and requirements might be used to indicate how the simulated mechanisms should be implemented. As a result, the capturer knows from an entity’s creation its entity type, the file’s location, the file name, and what other entities from the conceptual model were used in its specification. After a new file has been created, the capturer creates a provenance event notifying the provenance builder that the respective entity has been created. In addition, the provenance capturer then tracks the entity’s file to detect when it is edited. When a tracked file is edited, the capturer will send an event of the type “specifying <entity type>”, including the old and new versions of the file, to the provenance builder.

The Python utility library “simprovhelper” provides functions that extract information (like the script’s content, used libraries, their versions, etc.) from scripts, experiment specifications, and their outputs. When a modeler creates a simulation experiment from the command palette, then that new file will already call a function from the “simprovhelper” library to collect all information about the experiment’s execution and contain a comment explaining how to use that function. The modeler can then implement their simulation experiment as they are used to. Before running the experiment for the first time, they only have to add the simulation model’s path and the path to the simulation experiment’s output data as parameters to the library function (as stated in the explanation in the comment). Now, once the modeler executes their experiment, the capturer will detect this activity because of the library function and collect all information associated with running an experiment. Then, as part of the library function, the capturer sends the gathered information to the provenance builder as a provenance event “executing <entity type>”.

These two steps instrumenting Visual Studio Code with a provenance capturer are only necessary once for both of the following case studies and can also be used for future studies. As long as we continue to use Visual Studio Code and Python and the same provenance model, we can use this instrumentation and do not need to install or create a capturer repeatedly.

### mRNA Delivery Model

Understanding and exploiting the process of delivering mRNA into cells has been widely identified as one of the key challenges for developing vaccinations, protein replacement therapies, and treatments for genetic diseases [82–84]. One potential method involves the utilization of lipoplexes [85], i.e., small lipid spheres that shuttle into the cell and unpack the encapsulated mRNA.

In [30], the authors developed a simulation model that accurately predicts key features of the delivery process via lipoplexes, e.g., the likelihood of synthesizing a target protein in relation to the dose of lipoplexes placed in the cells’ environment. However, the simulation showed clustering in the absolute height of the target-protein expression curves (see Fig. 9, left), which was not observed in wet-lab experiments. The authors identified the use of a fixed amount of mRNA particles per lipoplexes, originating from the limitation of the simulation software, as a likely cause of this behavior.

In [31], we developed an efficient simulator for a variant of ML-Rules [80], i.e., ML-Rules 3. We used the lipoplex model implemented in ML-Rules 3 to showcase the features and the performance of the new simulator, as well as the impact of considering dynamic compartments on the simulation results. In the following, we will show how *SIMPROV* captured the provenance information while reimplementing the original model in ML-Rules 3, which allows the use of a varying number of mRNA per lipoplex.

#### Captured Provenance Graph

The captured provenance graph of the simulation study consists of 153 nodes and 342 edges, comprising all details about the simulation study’s entities, activities, and agents. This fine-grained provenance information allows for tracing and introspecting all study products throughout the simulation study, significantly contributing to the desired reproducibility. However, it also introduces a challenge in providing an overview of the different phases within a simulation study.

The reduced provenance graph, as generated by *SIMPROV*, provides an overview of the different phases by aggregating chains of the same provenance activities into a single activity, resulting in a more coarse-grained provenance graph (see Fig. 6).

**Fig 6.**
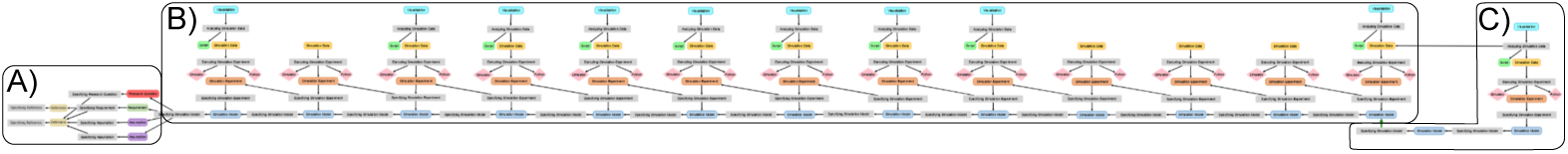
Reduced provenance graph showing the phases of the simulation study, including the A) the specification of the conceptual model, B) the step-wise re-implementation of the original model, and C) the adaptation of the simulation model and comparative visualization of the simulation results of both models.

From a birds-eye view, we can identify three phases: A) the specification of the conceptual model, B) the stepwise re-implementation of the original simulation model in ML-Rules, and C) the adaptation of the simulation model followed by comparative visualization of the simulation results of both models. In the following, we give an overview of each of the phases.

#### A) Conceptual Modeling

The simulation study started with the development of the conceptual model (see Fig. 7). For this, we began with the specification of references referring to a publication about the delivery process via lipoplexes and the source of the original simulation model.

**Fig 7.**
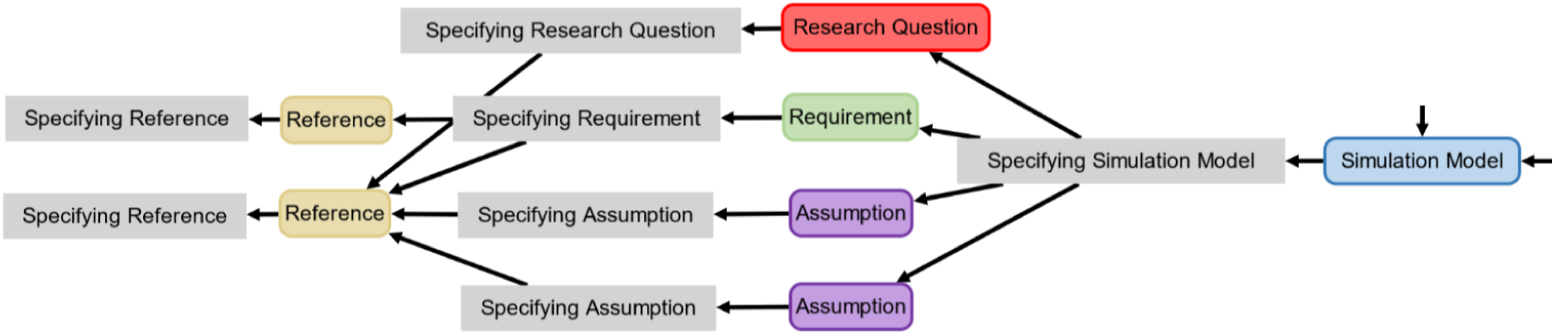
Overview of the conceptual modeling phase of the simulation study.

Based on these references, the research question was specified, i.e., re-implement the original simulation model and analyze the models’ behavior for varying amounts of mRNA particles per lipoplex. Further, we used the references to deduce crucial assumptions for the simulation model, i.e., the uniform distribution of all simulated particles and the stochastic behavior of the system. Finally, we specified the requirement that our simulation model be able to reproduce the results shown in the original publication.

From the modeler’s perspective, *SIMPROV* captured the provenance information of the entire conceptual modeling process of the simulation study. However, the modeler had to manually specify the relation between the different conceptual aspects as part of their specification, e.g., using a reference for specifying an assumption, as these relations could not be inferred.

#### B) Re-implementation of the Original Model

After developing the conceptual model, we started re-implementing the original simulation model following an incremental approach in ML-Rules (see Fig. 8). In each iteration, our simulation model was extended by another reaction rule, e.g., a rule to describe the attachment procedure of lipoplexes to the cell. The newly added reaction rule was tested, e.g., whether and how often the reaction rule fired and whether the dynamics of the reaction rule and the model appeared plausible. For this, an initial simulation experiment was developed and subsequently adapted to generate the necessary simulation data to be visualized. The iterations continued until all mechanisms of the original model were integrated as reaction rules and the original simulation data were reproduced (see Fig. 9 left).

**Fig 8.**
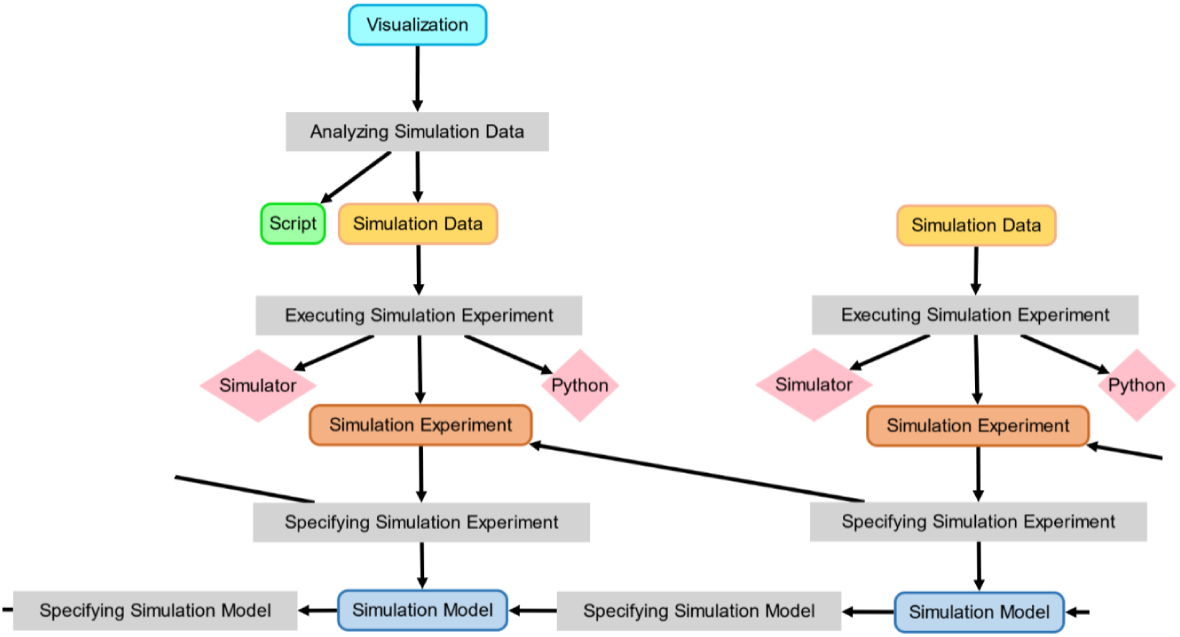
Overview of an iteration of the re-implementation phase of the simulation study. The modification on the right-hand side led to an error and an empty simulation data file that could not be visualized.

**Fig 9.**
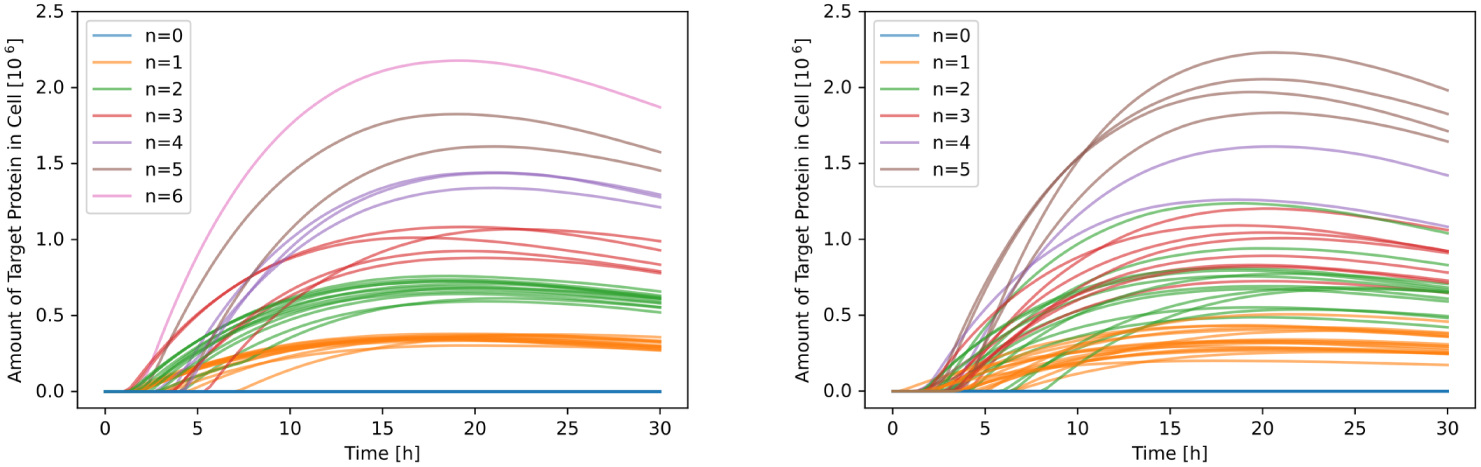
Comparative visualization of the simulation model for a fixed and varying amount of mRNA per lipoplex particle. The colors indicate the number of lipoplexes that unpack their mRNA in the cell. Left: Fixed amount of 350 mRNA per lipoplex. Right: Varying amount of mRNA per lipoplex.

In contrast to the specification of the conceptual aspects, all relevant provenance information from this phase is directly captured from Python scripts responsible for generating and visualizing the simulation data. Thus, capturing the provenance information of this phase integrates seamlessly into the modelers’ workflow.

#### C) Adaptation Of the Model

In the simulation study’s final phase, the model was adapted to support a varying amount of mRNA particles per lipoplex, which was not possible in the original model.

For this, we copied the ML-Rules model (of B) and removed the fixed initial solution from the specification. As a consequence, we now rely on the simulation experiment to dynamically generate an initial solution with a varying amount of mRNA, i.e., by sampling the diameters of the lipoplexes from a normal distribution and calculating the number of mRNA particles based on the equation given by [86], and inject it into the model specification before running the simulation. The resulting simulation data of the different models were visualized (see Fig. 9). In visualization, we see a significant reduction of the clustering in the different trajectories of our adapted simulation model.

Similar to the previous phase, all the provenance information from the simulation experiments and visualization of the results are directly captured. However, the cloning and adaptation of the original simulation model initially appear as a “Specifying Simulation Model” activity generating a new simulation model. The relation to the old simulation model cannot be inferred using the rules. Thus, the modeler manually added a dependency (green arrow) to make the relations between both models explicit.

### Wnt/*β* Catenin Model

The Wnt/*β* catenin signaling pathway is essential during embryonic development and adult tissue homeostasis [87]. One part of this pathway is the internalization of the LRP6 receptor. In the paper [18], four different mechanisms of internalization were modeled and compared to wet-lab data. The model-building process is documented in a provenance graph. We reimplemented the first two models (M1 and M2) in Tellurium to compare the manually drawn provenance graphs to the provenance graph produced by *SIMPROV*.

No additional changes were necessary to use Tellurium with *SIMPROV* since the already existing utility library for the modeler can be reused to report the provenance information from the simulation experiments to the provenance builder. Since the capturing process was described in detail in the previous section, we will only focus on comparing the two graphs in this section. In addition, we will show how a working environment for simulation can be adapted to work with *SIMPROV* supporting an automatic documentation of provenance.

#### Manually vs. Automatically Generated Provenance Graphs

Fig. 11 shows the manually drawn provenance graph and Fig. 12 the automatically generated *SIMPROV* graph. There are two main differences between the two graphs. First, model M1 is based on five references in the original provenance graph, while the *SIMPROV* graph also lists assumptions, requirements, and a research question. However, these additional nodes are optional. The second difference is in the generation of data.

**Fig 10.**
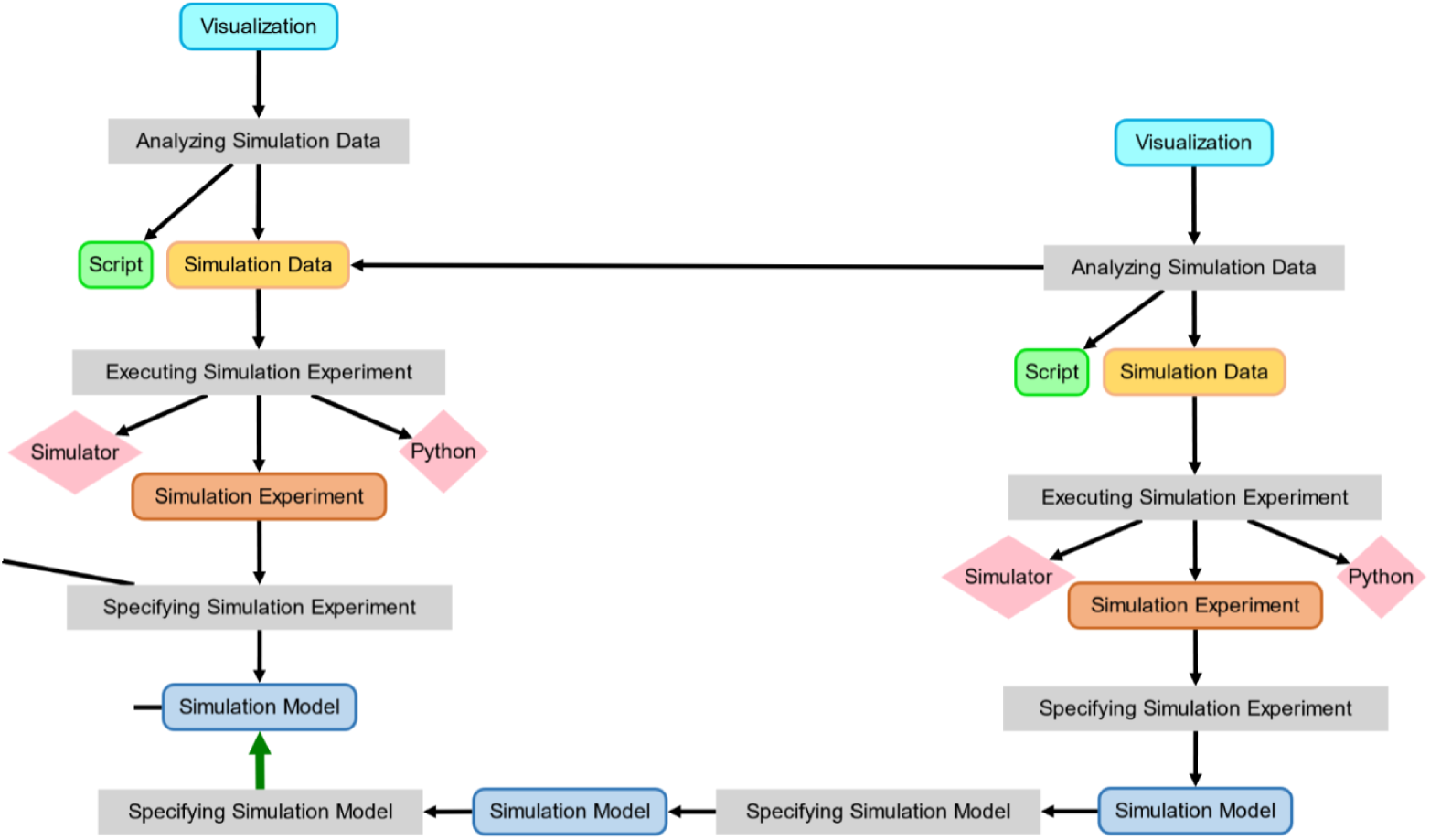
Adaptation of the simulation model and visualization of the different simulation results. The green arrow indicates a dependency manually added by the modeler to describe the relation between the re-implemented original model in ML-Rules 3 and the adapted simulation model in ML-Rules 3.

**Fig 11.**
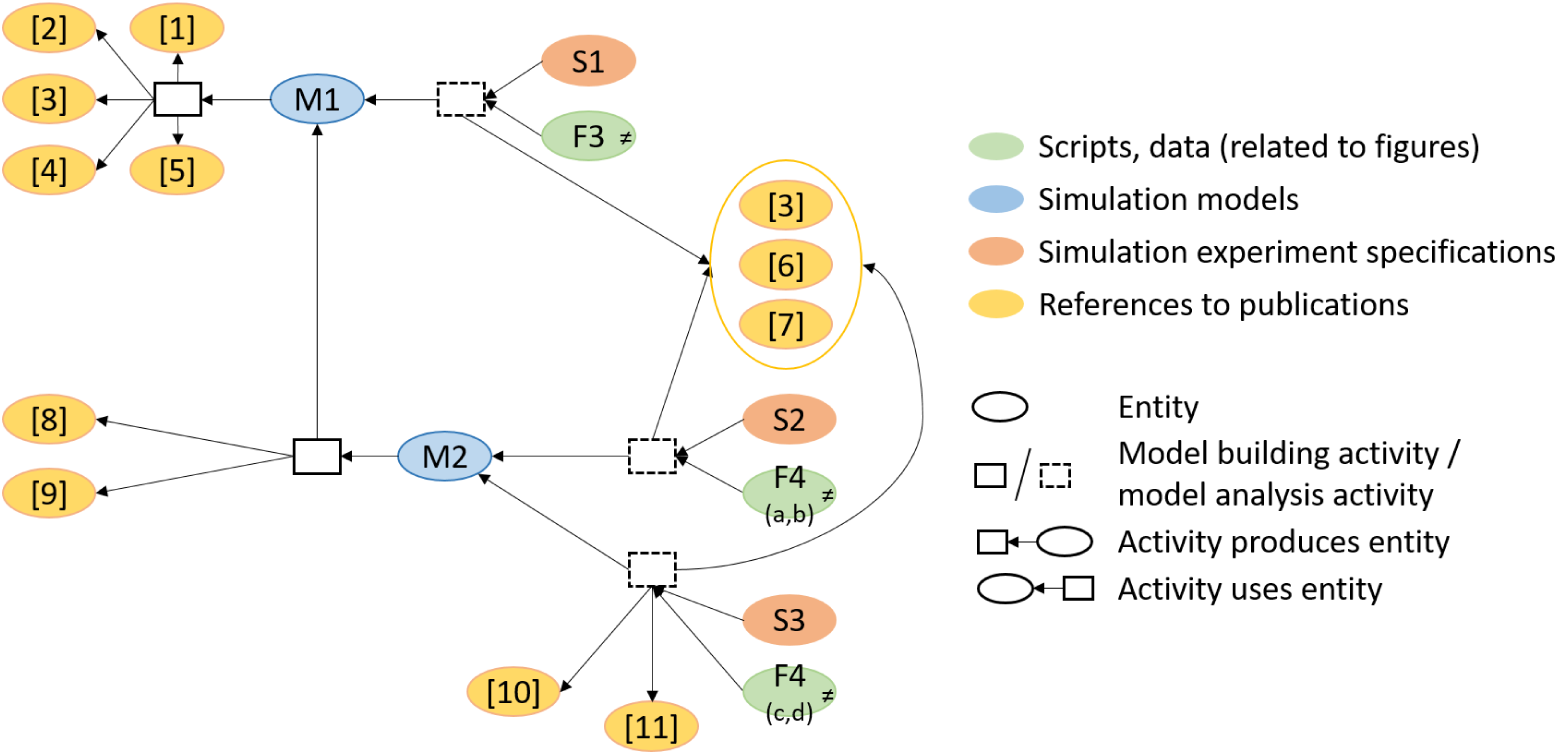
Original provenance graph from [18].

**Fig 12.**
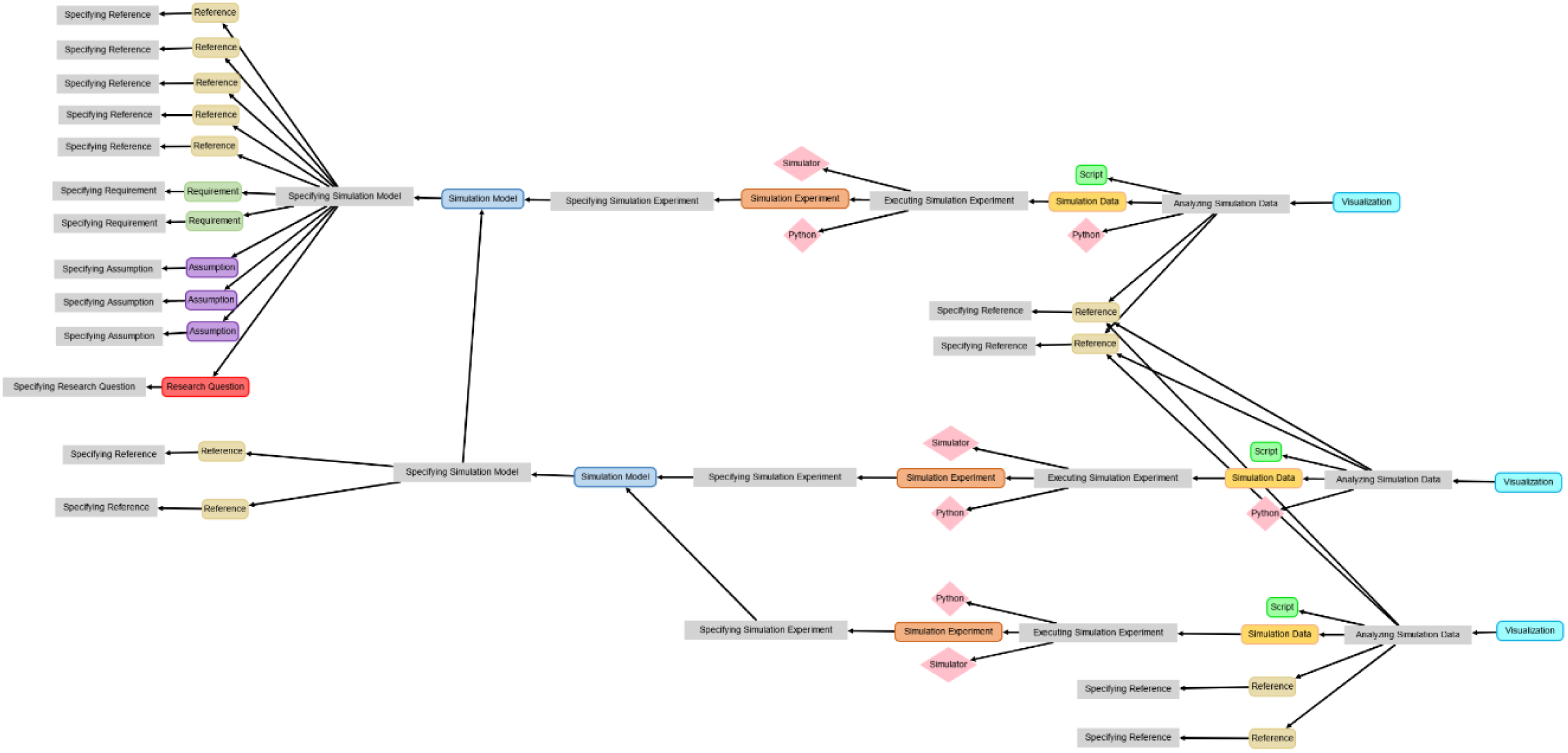
*SIMPROV* graph.

In the original graph, a simulation experiment and data (related to figures) are generated by an activity that uses the simulation model (and possibly other sources); this activity aggregates the specification and execution activities that are distinguished in the provenance graph produced by *SIMPROV*.

Despite these minor differences, both graphs are very similar in visualizing which references are used to build the different models and which data/figures are plotted from these models.

## Discussion

In developing *SIMPROV*, we set out to capture the provenance of entire simulation studies with minimal user involvement. Now, we will discuss the range of the automatically captured information, the degree of user interaction necessary, and the effort required to support new tools.

### What can be deduced from the modeler’s activities?

The provenance information that our approach can deduce highly depends on the implementations of the provenance capturers and how the modeler works. In the above case studies, *SIMPROV* with the Visual Studio Code capturer can fully automatically detect and record when a file is edited and when an experiment is executed by using plugin interfaces and wrapping library functions. When the modeler edits a file, both the old version of the file and the new one are recorded with the detected activity without any additional information from the modeler. Similarly, once the modeler executes a simulation experiment, the capturer records the activity, the executed experiment, and the resulting simulation data automatically. The provenance capturer can easily identify entity and activity types because the modeler chooses a command from Visual Studio’s command palette that creates a file specifically for the chosen entity type. And, finally, the commands prompt the modeler to choose which entities from the conceptual model are used to specify simulation models and experiments. So, all entities and activities described above can be recorded with *SIMPROV*.

### What has to be specified by interaction with the modeler?

However, to identify and record some of the activities we require the modeler to provide information that cannot be collected otherwise. The modeler has to tell the provenance capturer what entity types the files belong to and which entities from the conceptual model are used to specify simulation models and experiments. This paragraph will look into why we require these inputs from the modeler and how the information could be gained alternatively.

Since there are many different forms that the entities of simulation studies can take, no entity can always be identified unambiguously purely based on its form. Table 1 shows that some entities can occur in forms that are unique for the entity. For example, only simulation models can come in the form of simulation modeling formalisms.

However, they can just as well be specified using a general-purpose programming language that could also be used to specify simulation experiments or any other script. Therefore, the entity types of products generally cannot be discerned from the file type alone. For that reason, we require the modeler to name the entities explicitly by choosing the correct command (like in the case study above) or following a naming convention supported by the respective provenance capturer.

In addition, the modeler could use other tools to specify different products as long as each tool is equipped with a corresponding capturer. For example, they might use Visual Studio Code only to specify the simulation model and experiments. Then, they could use a wiki to document the conceptual model [40] which they instrumented with a capturer that sends events to the provenance builder whenever a wiki page is edited. In that case, their wiki structure would make the entity types of the respective products explicit, as well as the necessary metadata. Similarly, a documentation of the simulation study (e.g., Tellurium notebooks) could be instrumented with a capturer to automatically collect provenance (cf. instrumentation of Jupyter notebooks for experiment generation [88]). All in all, the modeler can use whatever tools they prefer, as long as the products’ entity types are made explicit and the tools are equipped with capturers.

But even if all the entities can be captured, it is difficult to automatically record which elements of the conceptual model were used to specify other products. For example, the modeler might read a description of a mechanism in a paper and then include that mechanism in their simulation model. Since there is no direct link from the simulation model file to the paper file, the dependency between the *specify simulation model* activity and the paper’s *information* entity has to be added manually by the modeler. Therefore, our Visual Studio Code capturer allows the modeler to select which elements from the conceptual model are used in an entity’s creation in our case study. Again, it might be possible to fully automatically capture the dependencies between activities and entities from the conceptual model in some cases. Take the example from above, where an experiment imports an assumption as a logic formula to check if the formula holds for a simulation model in a certain configuration. In this case, the assumption’s involvement in the experiment specification and execution can be inferred by inspecting the experiment file for imports. However, this only works for entities that are directly imported by another entity.

### How much implementation effort is needed?

To use *SIMPROV* in a simulation study, a modeler has to only start the *provenance builder* and use the software systems that already integrate *provenance capturers*. After a simulation study, a JSON representation of the provenance information [77] can be downloaded for further processing or publishing.

However, *SIMPROV* is also associated with a trade-off as additional effort is required to implement the *provenance capturers*, and to specify the *provenance patterns* for the *provenance builder*. Examples of this were shown in the case study for Visual Studio Code and Tellurium. However, since modelers tend to stick to their familiar modeling and analysis environment, once everything is implemented, no additional effort will be required for their future simulation studies.

### What insights do the captured provenance graphs provide?

Currently, *SIMPROV* improves the readability of the provenance graphs by aggregating chains of similar activities. The reduced provenance graphs convey information about the phases of conceptual modeling, simulation model specification, and model analysis via simulation experiments.

Yet, the captured provenance graph could still be further processed than in our current implementation to reveal even more insights into its simulation study. As shown in [29], different kinds of reductions may be applied on demand to generate distinct provenance views emphasizing different aspects of a conducted simulation study.

Moreover, it might also prove useful to capture and present the intentions behind the activities. For instance, experiments may be conducted for the purpose of calibration or validation [17]. Thus, the activity type *Execute Simulation Experiment* could be split into two activity types *Execute Validation Experiment* and *Execute Validation Experiment*. To identify these automatically, the types of entities used and generated by experiment executions could be held to account. The definitions of what types of entities are used and generated by experiments with which intentions are available in [22].

## Conclusion and Future Directions

A 2021 study showed that 49% of models in systems biology are not reproducible [24]. The main reasons were insufficient, incomplete, or ambiguous reporting. Thus, in this paper, we introduced *SIMPROV* as a lightweight method for automatically capturing provenance information during an entire simulation study with minimal user involvement. In the captured provenance graphs, not only the final products of the simulation study are reported, but also intermediate activities, products (including requirements and assumptions), and their interrelations. This explicit story of the simulation study can complement existing model documentation and facilitate the interpretation and reusability of results.

In our method, *provenance capturers* are responsible for detecting and collecting information about the modeler’s activity from the different software systems used in the simulation study. A *provenance builder* accepts this information, extracts, and validates provenance activity by declarative specifications of provenance patterns. Finally, the activity will be chained into a provenance graph, which can be further visually explored via a web interface. Further, we demonstrated how our software system could be applied and extended to illuminate the different phases of a simulation study using two-cell biological case studies.

The starting point for our approach has been that developing simulation models is an iterative and knowledge-intensive process involving the activities of domain-experts [89]. This also implied that some parts of the above process could not be automatically captured, e.g., which part of a publication was used in developing the simulation model. However, new methods challenge this traditional development of simulation models. In the last decade, significant advances have been made toward learning entire simulation models from data. The *reactionet lasso*, a structure learning approach, takes advantage of information-rich single-cell data, a tractable problem formulation, and partial knowledge about networks [90]. Other approaches are based on SINDy (Sparse Identification of Nonlinear Dynamics) [91] to estimate parsimonious biochemical reaction networks based on time series [92, 93]. Again, others combine information extraction from literature, clustering, simulation, and formal analysis to support an automated assembling, testing, and selection of context-specific models [94, 95]. All these approaches have in common that everything that is used to develop the simulation model has to be available and thus will also be accessible to the respective capturers. Consequently, these new developments will increase the part of simulation studies that becomes amenable to automatic capturing.

## Acknowledgements

This research was supported by the German Research Foundation (DFG), grant 320435134 (GrEASE).

## Software Availability

All mentioned software can be found at https://github.com/orgs/MosiSimProv/repositories. The complete documentation of the software can be found at https://simprov.readthedocs.io/en/latest/index.html. A software artifact containing all relevant files, including the specifications of the rules and provenance pattern, as well as provenance graphs of the case studies can be found at https://doi.org/10.5281/zenodo.14879929.

